# ChromBERT: Uncovering Chromatin State Motifs in the Human Genome Using a BERT-based Approach

**DOI:** 10.1101/2024.07.25.605219

**Authors:** Seohyun Lee, Junichi Sakatsume, Gina Miku Oba, Yuya Nagaoka, Che Lin, Chien-Yu Chen, Ryuichiro Nakato

## Abstract

Chromatin states, which are defined by specific combinations of histone post-translational modifications, are fundamental to gene regulation and cellular identity. Despite their importance, comprehensive patterns within chromatin state sequences, which could provide insights into key biological functions, remain largely unexplored. In this study, we introduce ChromBERT, a BERT-based model specifically designed to detect distinct chromatin state patterns as “motifs.” We pre-trained ChromBERT on 15-state chromatin annotations from 127 human cell and tissue types from the ROADMAP consortium. This pre-trained model can be fine-tuned for various downstream tasks, and obtained high-attention chromatin state patterns are extracted as motifs. To account for the variable-length nature of chromatin state motifs, ChromBERT uses Dynamic Time Warping to cluster similar motifs and identify meaningful representative patterns. In this study, we evaluated the performance of the model on several tasks, including binary and quantitative gene expression prediction, cell type classification, and three-dimensional genome feature classification. Our analyses yielded biologically grounded results and revealed the associated chromatin state motifs. This workflow facilitates the discovery of specific chromatin state patterns across different biological contexts and offers a new framework for exploring the dynamics of epigenomic states.

## Background

The understanding of chromatin organization is essential to reveal the complex mechanisms that govern gene regulation and function in the human genome[1], [2]. Chromatin states, defined by the combined features of histone modifications, including the involvement of DNA methylation and acetylation, are associated with specific functions such as gene activation, repression, or structural organization [3], [4]. For instance, active chromatin states—often marked by histone modifications such as H3K4me3 (trimethylation of lysine 4 on histone H3) and H3K27ac (acetylation of lysine 27 on histone H3) —are generally linked to gene activation and transcriptional initiation [5], [6]. These associations were further refined by Heintzman et al.[7], who distinguished promoter- and enhancer-specific marks, and Creyghton et al.[8], who identified H3K27ac as a marker of active regulatory elements. A deeper understanding of chromatin states and their genomic distribution has been facilitated by the chromatin state annotation tools such as ChromHMM [9] and Segway [10] coupled with the advent of chromatin immunoprecipitation followed by high-throughput sequencing (ChIP-seq) experiments [11], [12], [13]. Such annotation tools process genome-wide ChIP-seq datasets and systematically learn and characterize combinatorial patterns of histone modifications into distinct chromatin states, such as “active promoters,” “genic enhancers” and “heterochromatin.”

Large databases of epigenome information have been generated by international consortia, such as the ROADMAP project [14], the ENCODE project [15], and the International Human Epigenome Consortium (IHEC) [16]. For instance, the ROADMAP project characterized and annotated the chromatin state of 127 distinct human cell and tissue types. The chromatin state dataset can be used to perform various downstream analyses to decipher the hidden intricacies of the chromatin state landscape. As one such attempt, ChromDiff [17] compared a summarized set of chromatin states across different epigenomes (cell and tissue types), offering new insights into tissue-specific or age-dependent chromatin alterations. More recently, CSREP [18], a framework for deriving representative chromatin state maps for groups of samples, was developed by Vu *et al*. Utilizing an ensemble of multi-class logistic regression classifiers, this method enables the summarization of chromatin states for different groups of epigenomes with high resolution. Additionally, Epilogos aims to visualize the conserved chromatin states across different epigenomes based on the surprisal score measured from the rarity of the chromatin state distribution [19]. A similar method, established with a different metric, the conservation-associated activity score, was suggested by Libbrecht *et al* [20] as a novel annotation strategy that has the advantage of detecting more biologically functional regions.

Recently, deep learning approaches designed for natural language processing have shown great promise in deciphering complex biological patterns from large-scale genomic and epigenomic datasets. Transformer-based architectures have been especially effective in modeling sequential dependencies, leading to successful applications in biological contexts. For instance, DNABERT [21] has been developed for DNA sequence analysis, while Geneformer [22] and EPInformer [23] have demonstrated the potential of deep learning for modeling cellular states and integrating epigenomic data.

The availability of chromatin state annotations for a large dataset, recent advances in deep learning, and the precedent related studies have paved the way for the next important step: identifying patterns of chromatin states as ‘motifs’. A ‘motif’ refers to a recurring sequence that possesses a biological significance, often associated with specific functions such as binding sites in DNA sequences [24]. In the context of our study, we extend this concept to chromatin states to identify distinct patterns that could be key to understanding gene regulation. Instead of examining individual genomic loci separately, analyzing the patterns and combinations of chromatin states in relation to the surrounding regions, which we refer to as chromatin state motifs, could reveal the functional significance and regulatory roles of spatially coordinated epigenomic modifications that were previously unrecognized. By mapping these motifs, we may begin to decode the epigenetic language that potentially modulates gene expression patterns across different cell types and developmental stages. However, since chromatin state patterns are inherently dynamic and varying in length and structure, they may not be fully captured by traditional motif identification methods, such as k-mer-based algorithms, which often rely on simplistic representations of DNA sequence [25], [26]. This underscores the need for strategies that can accurately decipher the motifs of chromatin states and reveal their functional implications in gene regulation and chromatin organization.

In this study, we introduce ChromBERT, which is specifically tailored for the discovery of chromatin state motifs using the Bidirectional Encoder Representations from Transformers (BERT) model [27]. BERT has proven highly effective in natural language processing tasks and provides an important way to analyze sequential patterns in biological data [28]. By integrating the BERT model with the sequence alignment algorithm called Dynamic Time Warping (DTW) [29], ChromBERT effectively analyzes dynamic chromatin states, which extends the concept proposed by DNABERT for DNA sequence analysis [21]. ChromBERT is designed as a general-purpose model capable of learning and representing genome-wide chromatin state patterns across cell types. In this study, we evaluated the performance of ChromBERT on several tasks, including binary and quantitative gene expression prediction, cell type classification, and three-dimensional genome feature classification. Using the chromatin state annotations from 127 human cell and tissue types from the ROADMAP consortium, ChromBERT successfully identified chromatin state motifs, demonstrating its potential for predicting more complex genomic features. The source code of ChromBERT is available at https://github.com/caocao0525/ChromBERT.

## Results

### ChromBERT framework

Figure 1 provides an overview of the ChromBERT framework, illustrating the key components and workflow involved. We used the 15-state chromatin annotations defined by the ROADMAP project, which are based on a combination of five histone modifications. Initially, numerically annotated chromatin state labels ranging from 1 to 15 are converted into alphabetical codes ranging from A to O, resulting in the generation of a chromatin state sequence (Figure 1(a)). In the training phase, the chromatin state sequences are tokenized and fed into a BERT model for embedding (Figure 1(b)). We tokenized the chromatin state sequences into overlapping 4-mers by sliding one character, resulting in a large vocabulary size of 50,630 including standard special tokens. We also explored tokenization by sliding *N* characters at once (stride *N*, Supplementary Fig. S1). Since chromatin state sequences are more continuous than DNA sequences, we assumed that skipping multiple characters would not significantly affect prediction performance. This approach enables the use of longer input sequences and allows the model to learn long-range dependencies across regulatory regions.

**Figure 1.**
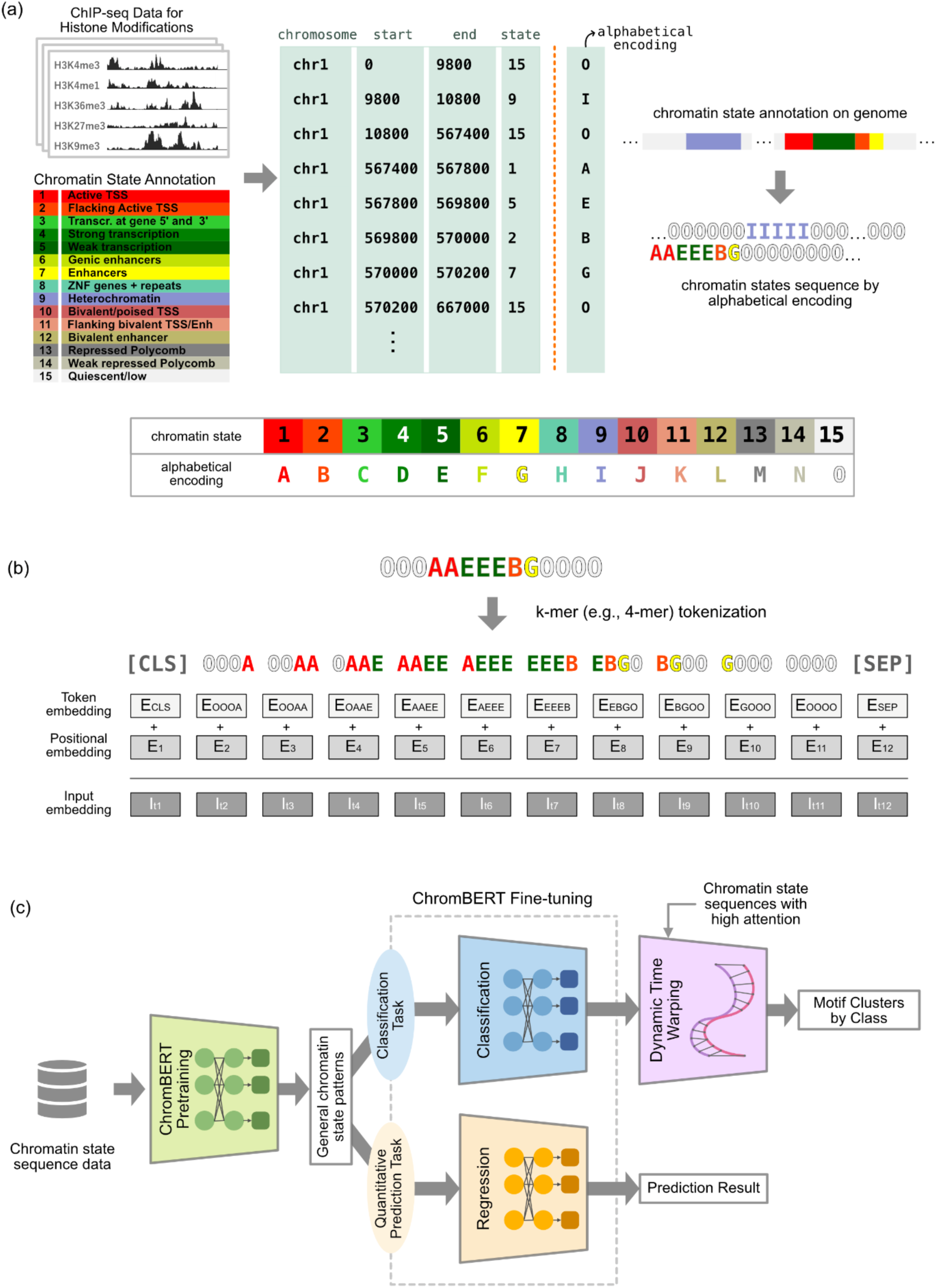
The ChromBERT workflow. (a) Conversion of numerical chromatin state annotations (from a BED file) into alphabetical encoding, followed by concatenation for generating a sequence of chromatin states (upper). The alphabetical encoding for each chromatin state (lower). (b) Creation of input embeddings for the BERT model through the combination of token and positional embeddings. (c) The comprehensive process of ChromBERT, from pretraining on general data to fine-tuning for task-specific applications.

The overall workflow of ChromBERT is captured in Figure 1(c), starting with the preprocessing of chromatin state sequence data. ChromBERT learns the general patterns of chromatin state placement through a pretraining stage. Following this, ChromBERT is fine-tuned for specific binary classification and quantitative regression tasks, such as promoter region classification and gene expression level prediction. As a downstream analysis of the classification task, chromatin state motifs specific to each task are extracted from the attention matrix values.

Motif discovery is a key downstream component of ChromBERT. Unlike DNA sequence motifs, chromatin state motifs lack well-defined reference libraries, making their identification and interpretation more challenging. Additionally, chromatin state sequences are inherently cell-type specific and dynamic. Even chromatin state patterns with similar regulatory functions can vary in length across genomic regions, reflecting both biological variability (e.g., differences in regulatory domain size) and technical variability (e.g., fluctuations in ChIP-seq signal-to-noise ratios). To address these challenges, ChromBERT employs a Dynamic Time Warping (DTW)-based approach to align and cluster motifs based on their structural similarity rather than exact sequence identity. This enables biologically meaningful motif discovery that accounts for the variability in chromatin state patterns. See the Methods section for further technical details.

### Pretraining ChromBERT

To learn generalizable representations of chromatin state sequences, ChromBERT was first pretrained using a masked token prediction objective on chromatin state sequences across 127 human cell and tissue types in the ROADMAP dataset. We pretrained ChromBERT in two ways for different use cases. First, ChromBERT was pretrained on promoter regions (from 2 kb upstream to 4 kb downstream of transcription start sites [TSSs]), as illustrated in Figure 2(a). These regions were chosen because they play a central role in gene regulation, integrating signals from nearby and distal cis-regulatory elements such as enhancers, silencers, and insulators [30]. Second, we pretrained the model on chromatin state sequences spanning the entire genome to support broader applications and facilitate flexible fine-tuning for other genomic regions or tasks. In this study, we primarily used the whole-genome–pretrained model.

**Figure 2.**
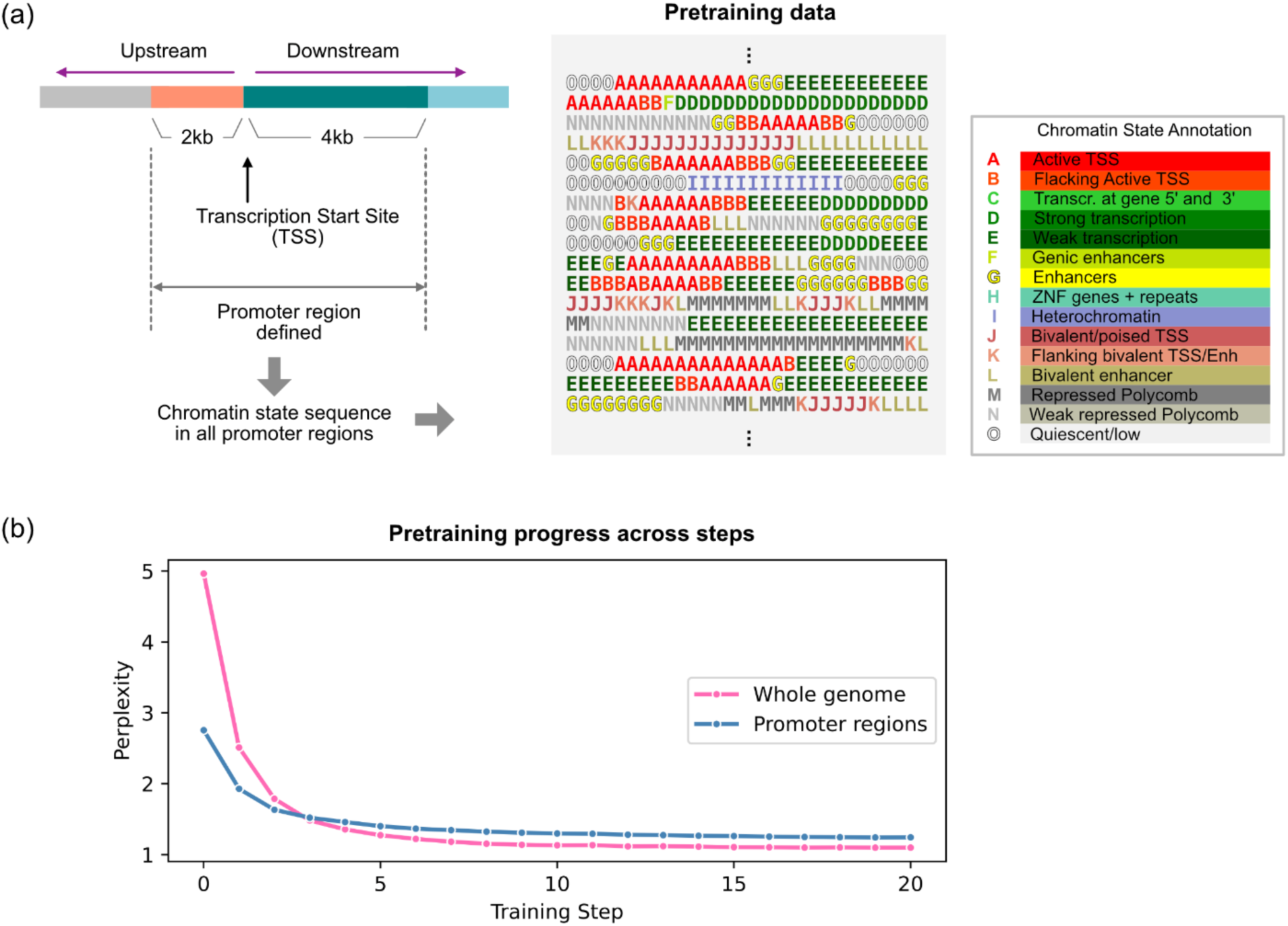
Overview of ChromBERT pretraining for promoter regions. (a) Definition of promoter regions used for pretraining. Chromatin state sequences were extracted from regions spanning 2 kb upstream to 4 kb downstream of TSSs across 127 cell types. The right panel shows an example of chromatin state sequences and a full legend of the 15-state annotation used in this study. The chromatin state annotation panel is from ROADMAP project [14] (b) Pretraining performance on chromatin state sequences from whole-genome and promoter-specific regions, measured by perplexity over training steps.

Throughout the pretraining process, the model’s perplexity decreased steadily, indicating successful learning of chromatin sequence structure (Figure 2(b)). For genome-wide pretraining, perplexity dropped from approximately 4.96 to 1.09. For promoter-specific pretraining, it decreased from approximately 2.75 to 1.24. The relatively low initial perplexity, particularly in promoter regions, likely reflects the repetitive and locally stable nature of chromatin state patterns near regulatory elements. See Methods section for further technical details.

### ChromBERT distinguishes between highly and lowly expressed genes based on chromatin state patterns of promoter-proximal regions

To evaluate the ability of ChromBERT to capture regulatory features, we first fine-tuned the model on chromatin state sequences around the TSSs to classify regions associated with genes of different expression levels. We used 57 cell types for which RNA-seq data are available in ROADMAP and evaluated the binary classification accuracy between highly expressed genes and genes with different expression levels. Genes were grouped based on their log-transformed expression values (RPKM). Genes with log-transformed RPKM values greater than 5 were labeled as ‘highly expressed,’ and we compared them with five groups: non-expressed genes and gene groups whose log-transformed RPKM values fell within 1–2, 2–3, 3–4, or 4–5. Figure 3(a) shows representative chromatin state sequences in promoter regions associated with highly expressed and non-expressed genes. Active states such as “active TSS” (“A”), “strong transcription” (“D”), “weak transcription” (“E”), and “genic enhancers” (“F”) frequently appear adjacent to highly expressed genes. In contrast, repressed states, such as Polycomb-associated states (“M” and “N”), dominate non-expressed regions.

**Figure 3.**
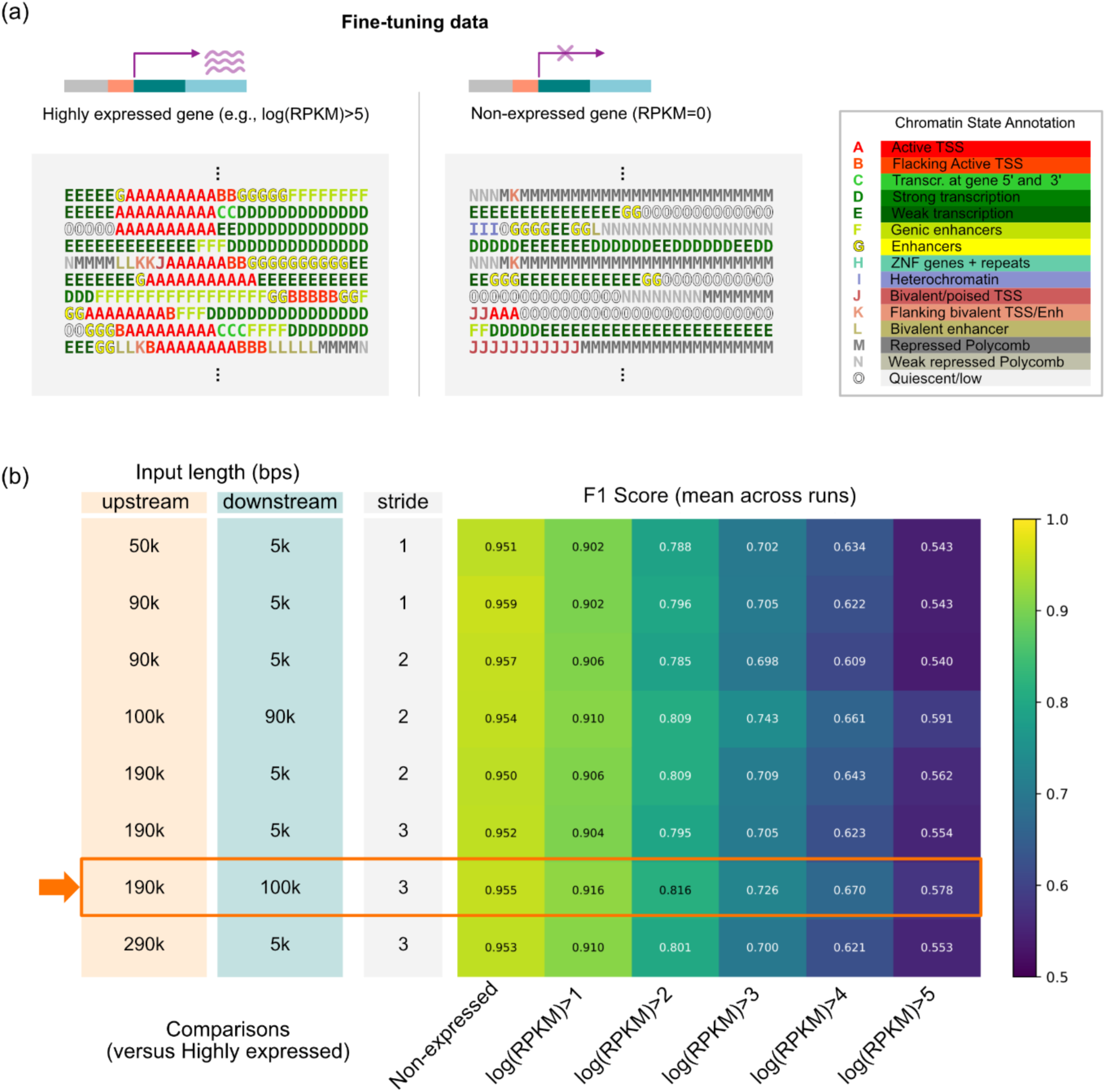
Fine-tuning ChromBERT on promoter regions for gene expression classification. (a) Example chromatin state sequences associated with highly expressed genes (e.g., log (RPKM) > 5, left) and non-expressed genes (RPKM = 0, right) used in fine-tuning. Color labels follow the 15-state annotation in ROADMAP (legend consistent with Figure 2). (b) Summary of the input-region configurations evaluated during fine-tuning. Upstream and downstream windows of varying lengths (50–290 kb upstream; 5–100 kb downstream) were tested using stride values of 1–3. The heatmap displays the mean F1 scores across runs for classification tasks comparing highly expressed genes with gene groups defined by different log-transformed RPKM thresholds. The configuration highlighted with an orange arrow corresponds to the up190k/down100k window. AUC and accuracy matrices for the same configurations are provided in Supplementary Fig. S2.

We next evaluated how the choice of input region around TSSs affects fine-tuning performance. We tested multiple combinations of upstream and downstream windows and summarized the resulting F1 scores in Figure 3(b) (AUC and accuracy shown in Supplementary Fig. S2). As expected, classification between highly expressed and non-expressed genes was consistently high across all input lengths, while comparisons against genes with higher expression levels were progressively more challenging and yielded lower scores. Across input configurations, we observed modest performance gains when upstream regions were extended from short (50 kb) to moderately long ranges (approximately 90–190 kb). Including downstream regions also improved performance, particularly when combined with longer upstream windows. The highest overall accuracy was obtained with the up190k/down100k configuration (orange arrow). These results indicate that promoter-proximal chromatin states over extended upstream and downstream regions contribute meaningful information for distinguishing transcriptional activity levels. Overall, genes with high and low expression were classified with high accuracy, suggesting that ChromBERT effectively captures the regulatory patterns underlying transcriptional activity.

### ChromBERT identifies promoter chromatin state motifs associated with high gene expression

We next examined how ChromBERT distributes attention across the input sequences to identify which regions contribute most strongly to the classification task. As shown in Figure 4(a) and 4(b), attention values were sharply concentrated around the TSS, indicating that ChromBERT consistently prioritizes promoter-proximal features even when provided with extended upstream and downstream contexts. Outside this central peak, we also observed smaller high-attention regions on both sides of the TSS, suggesting that additional regulatory patterns in nearby upstream and downstream regions contribute to the model’s decisions. These results highlight that promoter-proximal chromatin states serve as the primary informative signals for distinguishing expression levels, while adjacent regions provide supplementary contextual cues.

**Figure 4.**
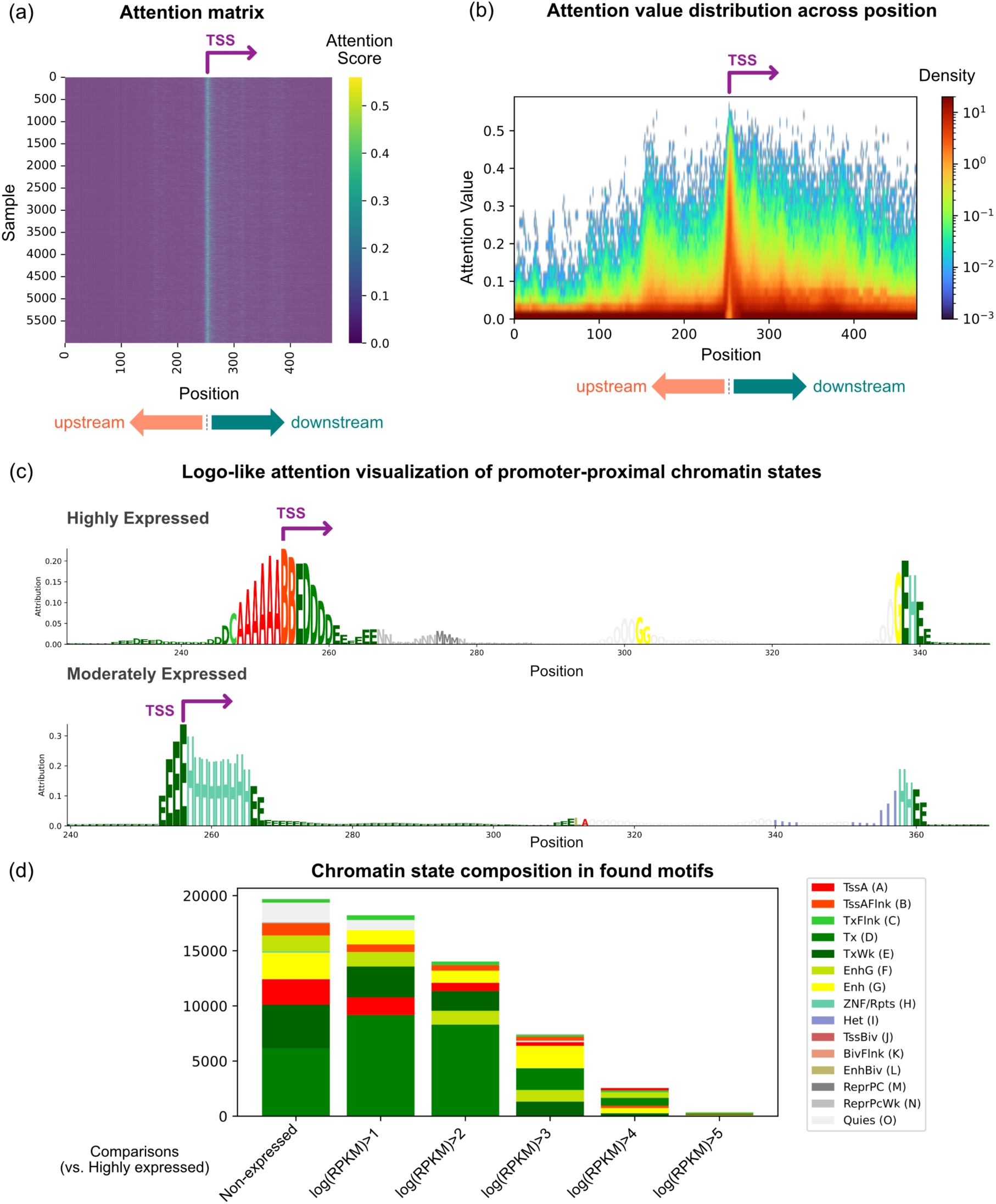
Attention-based interpretation of promoter-proximal chromatin states. (a) Attention matrix for representative samples from the fine-tuned model using a 100 kb upstream / 90 kb downstream input window around the TSS. Color values indicate normalized attention weights. (b) Positional density of attention values across upstream and downstream regions, aggregated over the evaluation set. (c) Logo-like visualization of promoter-proximal chromatin states. Bar heights are proportional to attention contributions. Examples are shown for a highly expressed and a moderately expressed gene. (d) Chromatin state composition of high-attention motifs for each comparison group, colored according to the 15-state annotation scheme.

To illustrate how the model focuses on individual promoter-proximal sequences, we visualized representative loci by plotting chromatin states with heights proportional to their attention values (Figure 4(c)). In the highly expressed example, the region surrounding the TSS shows high attention on promoter- and transcription-associated chromatin states, including “active TSS” (“A”), “flanking active TSS” (“B”), “strong transcription” (“D”), “weak transcription” (“E”), and with smaller contributions from “enhancers” (“G”). In a moderately expressed gene, attention around the TSS is lower overall and shifts toward transcribed and “ZNF genes + repeat” associated states (“H”), reflecting the chromatin context of this locus. Together with the global distribution of attention values (Figure 4(d)), these examples suggest that promoter- and transcription-associated states provide the primary signals for distinguishing expression levels, while nearby enhancer or repeat-associated states supply additional contextual information.

### DTW-based clustering reveals promoter chromatin motifs linked to high gene expression

The previous results suggested that chromatin state sequences tend to be persistent. Additionally, due to biological factors (e.g., different lengths of genes and enhancer regions) or technical factors (e.g., differences in read depth and signal-to-noise ratios across ChIP-seq samples), motif instances may vary in both character and length, even if they have the same biological functions. To summarize overarching patterns among the variable-length instances, we applied DTW and grouped similar chromatin state motifs into coherent families. This grouping uncovers recurring regulatory structures across genomic contexts post-fine-tuning, while still accounting for local flexibility in sequence patterns.

Initially, ChromBERT identifies motifs without imposing constraints on the merging. This approach captures a comprehensive set of chromatin state patterns that potentially signify the regions of interest, with selection based on statistical significance (p-value). ChromBERT then uses DTW to align the similar motifs in the set. Originating from the field of speech recognition, DTW is adept at accommodating variations in the tempo of analogous spoken words [31], making it particularly suitable for our purposes of aligning the different lengths of continuous sequences. DTW computes sequence similarities not by conventional point-to-point Euclidean distances but by minimizing the cumulative distances between motifs. Finally, the agglomerative clustering algorithm [32] is applied to cluster similar patterns into categories based on the sequence similarities, utilizing the hierarchical merging of data points based on their pairwise similarities.

The overall workflow of motif clustering is described in Figure 5(a). First, the pairwise DTW scores of motifs are calculated in both forward-forward and forward-reverse manners because the epigenomic patterns obtained by ChIP-seq do not distinguish between the forward and reverse strands. The lower DTW score representing higher similarities is adopted. Clustered motifs can be identified after determining the optimal number of clusters using a dendrogram. To visualize the clustering results, Figure 5(b) shows a UMAP projection of the motifs in a reduced-dimensional space [33]. This projection enables clearer assessment of how well the clusters are separated and reveals the overall structure of the motif space learned by ChromBERT. Figure 5(c) provides a cluster-wise summary that visualizes the composition and relative size of each cluster, allowing for intuitive interpretation of the distribution and prevalence of chromatin state motif patterns across samples. Together, these visualizations help convey both the diversity and consistency of motif types captured by ChromBERT. A line plot representation of all motif sequences by cluster, which preserves the full chromatin state transitions, is included in Supplementary Figure S3.

**Figure 5.**
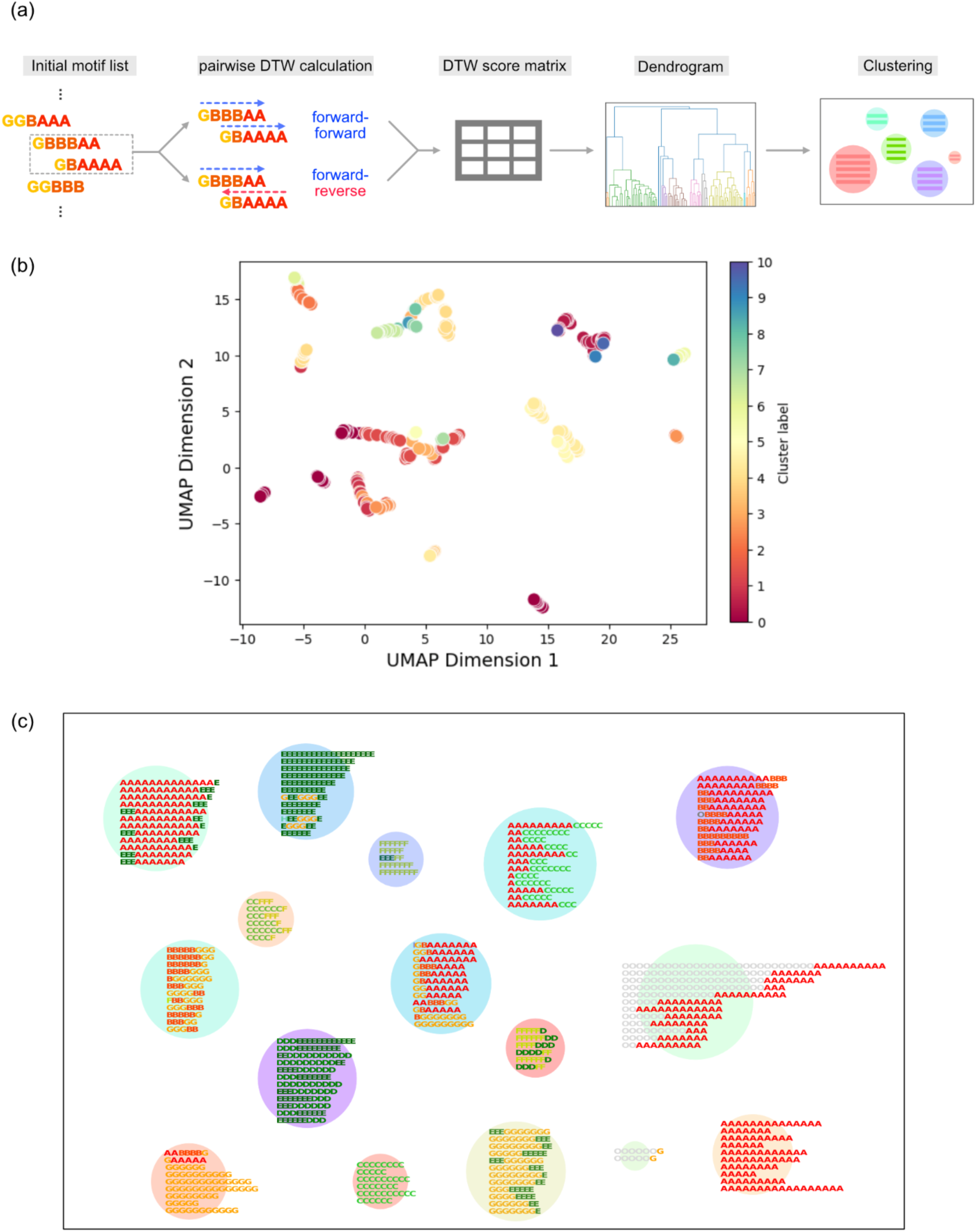
Chromatin state motif clustering with DTW. (a) Workflow of motif clustering using DTW. (b) UMAP representation of motifs extracted from the fine-tuned model using a 50 kb upstream / 5 kb downstream input window for the log(RPKM) > 5 vs. log(RPKM) > 1 classification task, clustered using agglomerative clustering. (c) Visualization of chromatin state motif clusters shown in (b), with each cluster’s size proportional to its number of elements.

To estimate the optimal number of clusters, we integrated several evaluative measures, including the dendrogram generated by the agglomerative clustering algorithm, to achieve a judicious balance between detail and overarching pattern recognition. In Figure 5(c), for instance, motifs with similar state transitions but different lengths, such as “BBBBGGG”, “BGGGGGG”, “GGGBBB”, were clustered into a single category, while “GGBAAAAAA,” which has a different transition, was clearly segregated into a separate category. Such transitions like “G-B-A” could represent enhancer regions immediately preceding active TSS regions, potentially acting as a herald for transcription initiation. This suggests a dynamic regulatory setup where enhancer activation might directly influence the onset of transcription at proximal promoter sites. Additionally, sequences such as “OOAAA” and “OOOOOAAA” in other clusters may reflect abrupt changes in chromatin states, possibly serving as precursors to promoter regions. Such patterns suggest the potential complexity of chromatin regulation captured by ChromBERT, and these motifs could provide interesting insights into the orchestrated events leading up to gene expression.

### ChromBERT predicts gene expression levels from promoter chromatin state sequences

To further expand ChromBERT’s utility, we adapted the model for regression tasks and attempted to predict gene expression levels quantitatively from promoter-proximal chromatin state sequences. The model was fine-tuned using log-transformed RPKM values associated with each gene.

We tested multiple combinations of upstream and downstream input lengths around TSSs (Figure 6a). Consistent with the binary classification, performance improved when both upstream and downstream regions were extended. The best configuration (100 kb upstream and 90 kb downstream) achieved a Pearson correlation of 0.791 between predicted and observed log-transformed RPKM values (Figure 6(b)), indicating that ChromBERT can quantitatively predict gene expression levels from chromatin state sequences. As illustrated in Supplementary Fig. S5, the categorical nature of chromatin state data leads to cases where similar sequence patterns exhibit different expression levels, which may constrain the maximum attainable correlation.

**Figure 6.**
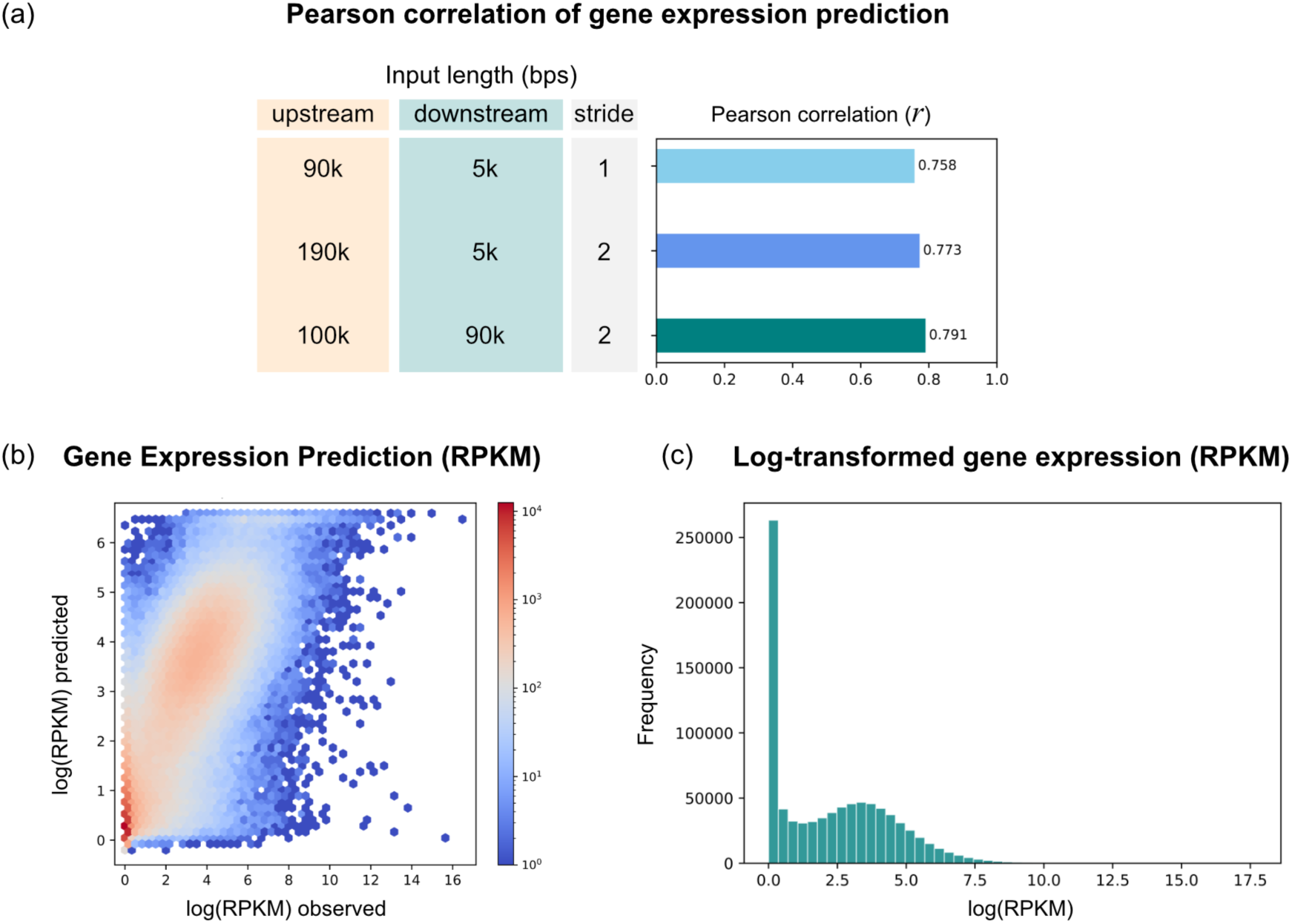
(a) Pearson correlation between predicted and observed log-transformed RPKM values for models trained with different upstream and downstream input windows around the TSS. (b) Hexbin plot showing predicted versus observed log(RPKM) values for the model using the 100 kb upstream / 90 kb downstream configuration. (c) Distribution of log-transformed RPKM values used for training and evaluation.

The gene expression density plot (Figure 6(c)) showed that most genes have log-transformed RPKM values below approximately 6, and ChromBERT could accurately estimate their expression levels within this range. In contrast, the model exhibited limitations for genes with log-transformed RPKM values exceeding 10, possibly due to the small number of such genes in the training data. Increasing the amount of training data for these genes would mitigate the imbalance and likely improve prediction accuracy.

### ChromBERT identifies cell-type–specific chromatin state signatures

To further evaluate the effectiveness of ChromBERT, we examined whether chromatin state sequences contain recognizable signatures of different cell-type groups and whether the model can classify them. We focused on cis-regulatory module (CRM) regions, which are enriched for regulatory activity. CRM regions were obtained from the SCREEN database [34], and chromatin state sequences were extracted from 20-kb windows centered on each CRM site. Cell-type group annotations followed the ROADMAP classification (“ESC,” “iPSC,” “ES-derived,” “blood-Tcell,” “HSC-Bcell,” “Brain,” “Muscle,” “Heart,” and “Smooth Muscle”), and we fine-tuned ChromBERT for binary classification across all pairwise combinations (Figure 7a).

**Figure 7.**
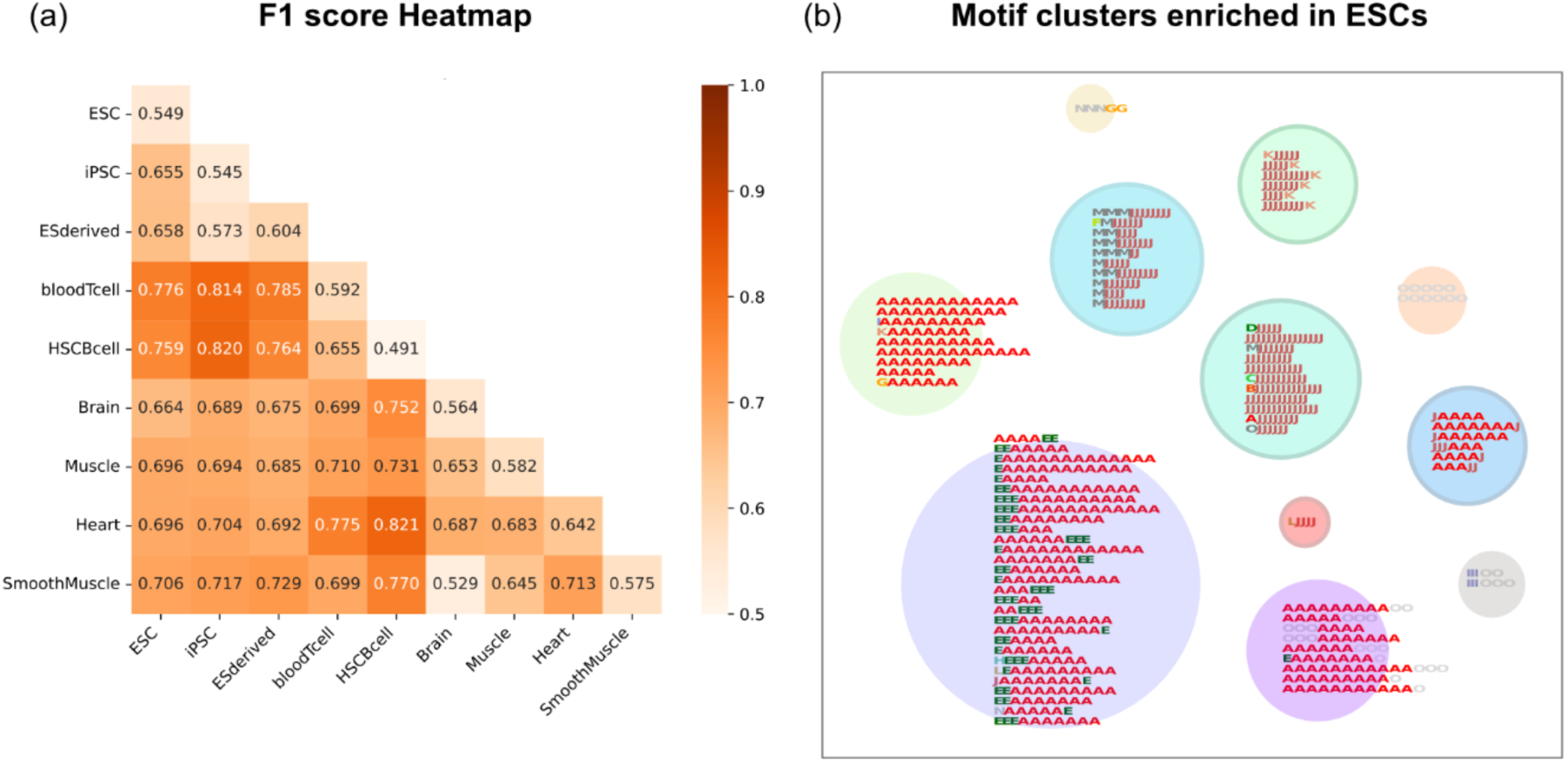
Cell-type classification and motif analysis across CRM regions. (a) F1 score heatmap for pairwise binary classifications of cell-type groups. Chromatin state sequences were extracted from 20-kb windows centered on CRMs obtained from the SCREEN database, and ChromBERT was fine-tuned for each pairwise comparison following the ROADMAP cell-type group definitions. (b) Motif clusters enriched in ESCs relative to T-cells, obtained from DTW-based clustering of promoter-proximal chromatin state motifs. Circle sizes are proportional to the number of motifs in each cluster.

The resulting accuracy distribution closely reflected known biological relationships among cell groups. For example, ESCs and iPSCs are highly similar in their epigenomic profiles and accordingly were difficult to distinguish (0.655). In contrast, ESCs and T cells showed more distinct chromatin patterns and were classified with higher accuracy (0.776). These results suggest that each cell group carries characteristic epigenomic signatures. One exception was the Brain vs. Smooth Muscle comparison, which had low accuracy (0.529). One possible reason is a problem with the data quality. According to the ROADMAP QC summary, multiple histone-modification ChIP-seq samples for these two cell groups were flagged as low quality[14]. These findings are consistent with those from the TSS-based gene-expression classification task (see Supplementary Fig. S4).

We next sought to identify motifs contributing to cell-type specificity. Figure 7(b) shows motif clusters enriched in ESCs when compared with T-cells. Interestingly, many ESC-specific motifs contain the character “J,” corresponding to the “bivalent/poised TSS” state, a hallmark of pluripotent stem cells [35], [36]. Motifs containing “J” appeared in in ESCs and iPSCs but not in differentiated cell types. A complete list of identified motifs is provided in Supplementary Table S1. These findings demonstrate that ChromBERT can reveal biologically meaningful, cell-type-specific motifs.

However, ChromBERT also identified motifs that appeared cell-type-unspecific (e.g., AAAAA). This observation suggests that differences between cell types may arise not only from biological distinctions but also from technical variability, such as differences in sample preparation or sequencing depth among cell groups.

### ChromBERT captures chromatin state signatures of 3D genome compartments

Lastly, we evaluated whether ChromBERT could capture features associated with three-dimensional (3D) genome organization. Although the relationship between the epigenome and 3D genome structure is still not well understood, if ChromBERT can accurately predict certain 3D genome features, it would suggest a strong correspondence between epigenomic states and higher-order genome organization. Using *in situ* Hi-C data from five cell types (GM12878, HMEC, HUVEC, IMR90, and K562) at 25-kb resolution obtained from Rao *et al.* [37], we first examined whether chromatin state sequences could distinguish A/B compartments (Figure 8(a)). See the Methods section for details on the Hi-C analysis. Compartments were defined using the first principal component (PC1) of the observed/expected contact matrix, where positive and negative PC1 values correspond to compartment A (generally active) and compartment B (generally inactive), respectively [38]. We additionally defined “strong compartment A” and “strong compartment B” as genomic bins with extreme PC1 values. As expected from their distinct chromatin compositions, ChromBERT accurately classified compartments A and B, with even higher performance for the strong A/B subsets where epigenomic differences were more pronounced.

**Figure 8.**
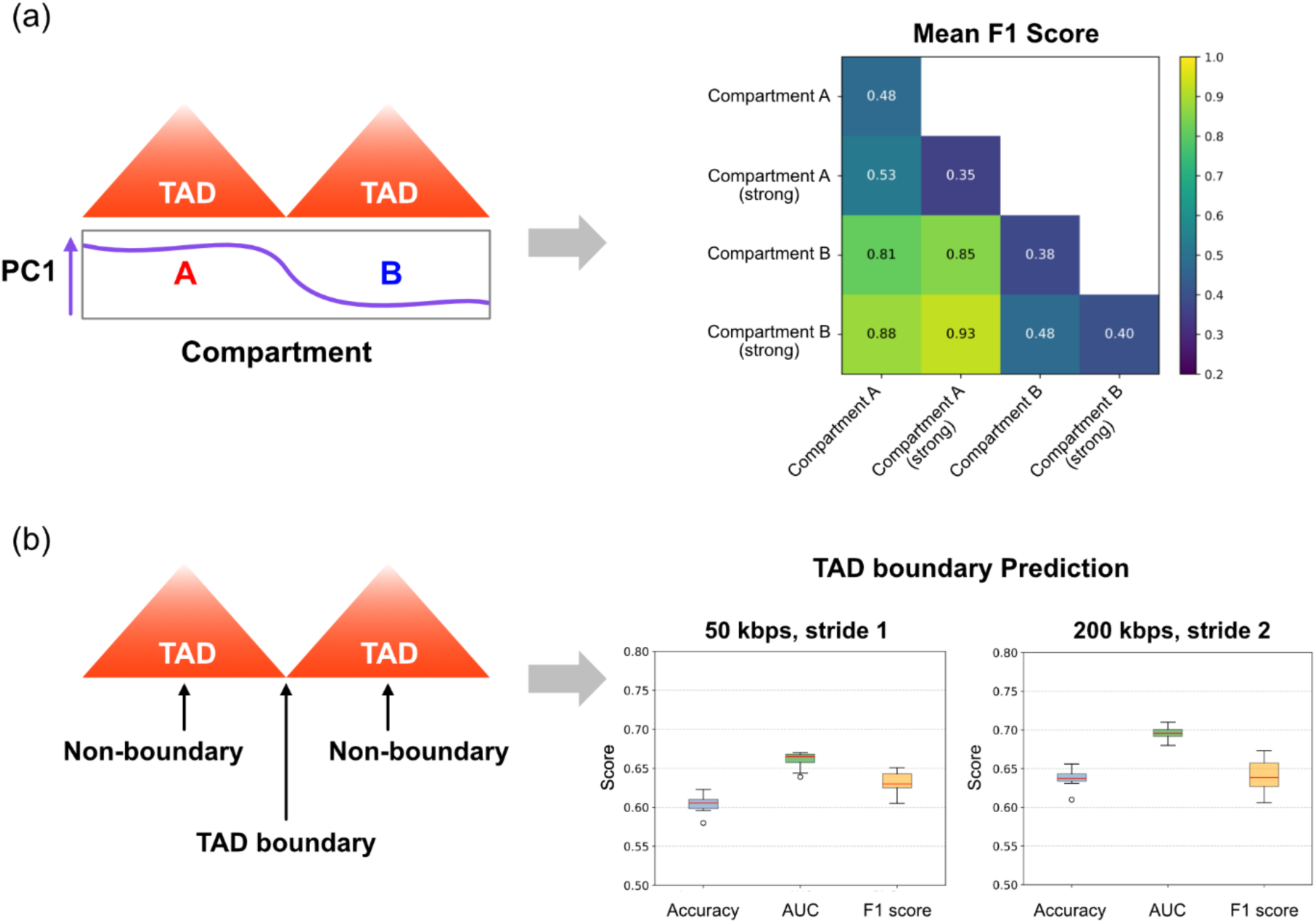
Classification of 3D genome features using ChromBERT. (a) Mean F1 scores for pairwise classification of A/B compartments and strong A/B compartments. Chromatin state sequences were extracted from genomic bins defined by the first principal component (PC1) of the observed/expected Hi-C contact matrix (25-kb resolution). (b) Classification performance for TAD boundary versus non-boundary regions. TAD boundaries were identified using insulation-score–based calling, and chromatin state sequences around boundary and non-boundary sites were used to train and evaluate ChromBERT with different input window sizes (50 kb and 200 kb).

We next attempted to classify topologically associating domain (TAD) boundaries versus non-TAD boundary regions (Figure 8(b)). TAD boundaries were identified from Hi-C contact maps following standard insulation-score–based procedures [39]. These regions are known to be enriched for the architectural protein CTCF and for housekeeping genes [40], suggesting that characteristic chromatin states might help distinguish them. However, unlike the compartment task, classification accuracy for TAD boundary versus non-boundary regions was modest (F1 < 0.7), and extending the input window to 200 kb resulted in only limited improvement. Several factors may contribute to this reduced performance. The number of TAD boundaries is much smaller than the number of compartment bins or gene-associated regions, and only five cell types were available, limiting the training data. Boundary calling also varies across computational tools, introducing technical variability. Moreover, both TAD boundary and non-boundary regions can contain mixtures of active and inactive chromatin states, reducing separability, and compartment information was not incorporated in this analysis.

Overall, these results indicate that large-scale features such as A/B compartments show strong correspondence with chromatin state patterns, whereas finer-scale structures like TAD boundaries are more difficult to predict from epigenomic sequences alone. Nevertheless, ChromBERT provides a useful framework for investigating how local chromatin states relate to 3D genome organization.

## Discussion

ChromBERT is a Transformer-based architecture specifically designed for chromatin state sequences. Unlike DNA sequence models such as DNABERT, which operate on static nucleotide sequences, ChromBERT learns from epigenomic annotations that are cell-type–specific and inherently variable in length. While recent deep learning approaches such as Geneformer [22] and EPInformer [23] have modeled regulatory states or integrated multi-omics signals, ChromBERT provides a complementary perspective by learning sequential patterns directly from chromatin state annotation data. By leveraging self-attention mechanisms, ChromBERT learns recurrent chromatin state motifs directly from sequence-level data, capturing structural patterns that are not the focus of annotation summarization tools like CSREP [18], alignment-based methods such as EpiAlign [41], or the conservation-based approach like Epilogos and the conservation-associated activity score [20]. This sequence-level viewpoint offers a data-driven framework for studying local epigenomic organization and its relationship to transcriptional regulation.

Across multiple downstream tasks, including gene expression prediction, cell-type classification, and the prediction of selected 3D genome features, ChromBERT demonstrated that chromatin state sequences encode biologically meaningful patterns. For example, promoter-proximal chromatin states enabled accurate binary and quantitative gene expression prediction, and cell-type–specific CRM regions yielded signatures consistent with known epigenomic programs (e.g., enrichment of the bivalent/poised TSS state “J” in pluripotent cells [35], [36]). Chromatin states also captured large-scale 3D genome organization: A/B compartments were readily classified, with stronger compartment regions showing particularly high separability. These results support the idea that chromatin state annotations retain regulatory information that can be learned directly at the sequence level.

A notable strength of ChromBERT is its scalability with respect to pretraining data. Because chromatin state sequences differ across cell types, the number of available training samples naturally increases as more cell types are incorporated. We also pretrained ChromBERT using chromatin annotation data from the IHEC project [16], which provides 1,699 epigenome annotations based on an 18-state model. Once the IHEC database becomes publicly available, the corresponding pretrained model will also be released. However, increased data volume also magnifies technical variation. Public ChIP-seq datasets vary in quality, and including samples with weak signal-to-noise ratios may introduce noise during pretraining. Similarly, many cell types are represented by multiple donors or slightly modified conditions, potentially leading to imbalanced datasets. Careful data curation and quality filtering will therefore be essential for maximizing model performance.

From a computational perspective, chromatin state modeling introduces unique challenges. DNABERT’s nucleotide alphabet consists of only four bases, resulting in manageable k-mer vocabularies (e.g., 4,101 (6-mers)). In contrast, ChromBERT must represent 15 chromatin states, and the resulting k-mer vocabularies expand sharply in absolute size: 3,379 (3-mers), 50,629 (4-mers), 759,379 (5-mers), and over 11 million (6-mers). This pronounced difference in vocabulary size is computationally challenging and can lead GPU memory limitations when attempting to pretrain with more than 5-mers, even with a single sample set. This effectively limited us to using 4-mers for the pre-training step. The vocabulary complexity becomes even more substantial when using the 18-state chromatin annotation system from the IHEC dataset.

Additionally, the perplexity of the model (a metric that quantifies how well a model predicts a sample) showed a decreasing trend from approximately 4 to 1 over iterations during the pretraining for both promoter regions and broader genic regions. Interestingly, the perplexity was already relatively low at the beginning of training (approx. 4). This suggests that the model is successfully predicting the masked tokens in the pretraining, even in the early stages. This initially low perplexity can be attributed to several factors. First, the stable, long-lasting nature of chromatin states and their functional bias likely lead to more predictable sequence patterns, easing the model’s predictions. Second, our use of 4-mer tokenization further simplified the task for the model, as it only needs to predict patterns within a more limited context compared to larger k-mers.

ChromBERT achieved strong predictive performance across multiple tasks. In gene expression regression, the model reached a Pearson’s *r* of 0.79 and captured promoter-proximal regulatory patterns that reflect relative transcriptional activity. Extending the promoter window improved performance up to a point, indicating that distal regulatory features contribute additional information. Chromatin state sequences also facilitated the discovery of recurring regulatory motifs, as their categorical representation integrates multiple histone marks and enables long effective input windows.

Like other BERT-based architectures, ChromBERT is subject to inherent input-length constraints, limiting the maximum sequence to 510 tokens. However, the locally stable and consecutive nature of chromatin states allows this constraint to be partially mitigated through stride-based tokenization, which enables the model to incorporate a broader genomic context while maintaining meaningful patterns. In addition, while 4-mer tokenization during pretraining does not directly restrict the motif length, its contextual granularity may bias the model toward learning patterns on a similar scale. To detect longer motifs, it may be useful to modify the sequence length constraints or revise the tokenization process to better accommodate the unique properties of chromatin state annotation data. Such efforts could further improve ChromBERT’s utility in decoding the complex, hierarchical nature of epigenomic regulation.

Unlike previous methods that use regression-based models to directly predict gene expression from raw histone-modification signals, chromatin state data does not use peak-intensity information. This could result in reduced performance for highly localized predictions. However, chromatin states offer a simplified sequence representation that integrates multiple histone marks, enabling much longer input sequences and allowing the model to focus on the length and combinatorial patterns of state transitions. This representation enables the detection of recurring regulatory patterns across samples and facilitates the discovery of sequence-level chromatin motifs that generalize across cell types. Furthermore, this framework is not limited to gene expression prediction. It can also serve as a foundation for many downstream tasks, such as cell type classification and 3D genome-related analyses.

Performance for TAD-boundary prediction was modest, likely reflecting both technical and biological factors. TAD boundaries are relatively few per cell type, vary by caller, and can contain mixtures of active and inactive chromatin states, reducing separability. Incorporating additional modalities, such as CTCF binding [42] or more finely resolved regulatory annotations, may improve the model’s ability to identify finer-scale 3D architectural features as multi-omics datasets expand.

Future work may explore model architectures with extended context, hierarchical tokenization strategies, or dynamically learned segmentation to better capture long-range epigenomic organization. Improved curation of input datasets, inclusion of additional cell types, and integration of complementary epigenomic modalities may further enhance ChromBERT’s generalizability.

In summary, ChromBERT provides a scalable, interpretable framework for learning sequence-level patterns within chromatin state annotations. By capturing recurrent motifs, promoter activity signatures, and compartment-level structures, ChromBERT illustrates how language-model approaches can decode the structured complexity of the epigenome and link local chromatin organization to diverse functional genomic outcomes.

## Methods

### 1. Data collection

The chromatin annotation BED files are downloaded from the ROADMAP [14] project, which includes 127 distinct epigenomes annotated with 15 different chromatin states at 200-bp intervals using ChromHMM [9] for genome build hg19. The 15 different chromatin states are 1. Active Transcription Start Site (TSS), 2. Flanking Active TSS, 3. Transcription at Gene 5’ to 3’ Ends, 4. Strong Transcription, 5. Weak Transcription, 6. Genic Enhancers, 7. Enhancers, 8. Zinc Finger Protein (ZNF) Genes & Repeats, 9. Heterochromatin, 10. Bivalent/Poised TSS, 11. Flanking Bivalent TSS/Enhancer, 12. Bivalent Enhancer, 13. Repressed Polycomb, 14. Weak Repressed Polycomb, and 15. Quiescent and low signal. The chromatin annotation BED files consist of four columns: chromosome number, start position, end position, and the number corresponding to the annotated state.

### 2. Data preprocessing

To prepare the data for natural language processing techniques, we first converted the chromatin-annotated data, originally represented by numbers 1 to 15, into corresponding alphabetic characters A to O. This transformation was necessary because natural language processing cannot be directly applied to sequences of numerical chromatin states. After changing the chromatin state annotations to alphabetic symbols, we condensed the genomic positions by a factor of 200 since the original annotations were conducted at 200-bp intervals. Consequently, we generated a continuous string of chromatin states, with each alphabet character (A to O) representing the corresponding chromatin state (1 to 15) for a 200-bp segment. Additionally, to focus our analysis on the relevant genomic regions and to exclude potential biases from telomeres, we removed the initial and final 10,000 bps from each chromosome, as these regions generally represent the extreme ends and may not accurately reflect the overall chromatin states. In this manner, all the chromatin annotation data for 127 epigenomes were processed chromosome-wise. Since we adapted BERT architecture, the processed chromatin annotation data were randomly cut to ensure that the length of the data did not exceed 510 but was longer than 5, following their data preparation method. Subsequently, the chromatin state sequences were tokenized into 3-mers, 4-mers, 5-mers, and 6-mers. Any sequences shorter than the length of each k-mer were not included in the dataset.

### 3. Pretraining

The BERT architecture utilized in ChromBERT was pre-trained on chromatin annotation data using datasets from 127 different cell types provided by the ROADMAP project for the 15 chromatin state case. The chromatin annotation sequences for the genic regions of these epigenomes were preprocessed as described in the data processing section, concatenated, and then tokenized using k-merization. Unlike DNABERT, we focused on testing 3-mers and 4-mers only, as 5-mers and 6-mers have a significantly larger vocabulary size (759,380 and 11,390,630, respectively), making them impractical to run. The vocabulary sizes for 3-mers and 4-mers were 3,379 and 50,630, respectively. Although both 3-mers and 4-mers demonstrated sufficiently low perplexity, we opted for 4-mers because they comparably capture more complex patterns more effectively, enhancing our analysis. The transformer architecture used for pretraining remained consistent with DNABERT, featuring an intermediate layer size of 3,072 and a hidden layer size of 768. To ensure smooth training without encountering out-of-memory errors, we adjusted the number of train and evaluation batches per GPU to 5 and 3, respectively. For other training conditions, we followed the same strategy as DNABERT, including a linear increase in the learning rate from 0 to 4e-4 during the warm-up phase. The pretraining process was conducted on a GPU machine equipped with two Nvidia GeForce RTX 2080Ti cards, and it took approximately three days to complete.

For the IHEC dataset, chromatin state annotations consist of 18 states (A–R) defined at 200-bp resolution. These sequences were converted into 4-mers to construct the pretraining corpus, yielding approximately 41 GB of data after removing tokens composed entirely of the quiescent state R. Because the full dataset could not be loaded into memory at once, we adopted a sequential pretraining strategy. The data were first randomly shuffled to distribute cell-line–specific patterns uniformly, then split into chunks of 500,000 lines. Each chunk was used in succession, with model weights carried over between chunks, enabling efficient training while avoiding overfitting to contiguous regions from individual cell lines. Pretraining was performed using a DNA-based Transformer architecture with 4-mer tokenization and a vocabulary of 104,980 tokens, representing all valid 4-mers from the 18-state system (excluding the RRRR token) together with the standard special tokens ([UNK], [PAD], [CLS], [SEP], [MASK]).

During pretraining, we used a masked language modeling (MLM) objective, following the standard BERT pretraining strategy. Specifically, 15% of tokens were randomly masked, as in DNABERT, and the model was trained to predict these masked tokens using surrounding context. Perplexity was used as the primary evaluation metric to assess learning performance. While chromatin state sequences exhibit structured dependencies, the self-attention mechanism enables ChromBERT to effectively capture both local and long-range relationships. Gradient accumulation and a carefully chosen batch size were used to efficiently manage GPU memory and allow training on large datasets. Model weights generated after each chunk were saved and used as initialization for subsequent chunks, progressively refining the model across the dataset. The training was conducted on a system equipped with an AMD EPYC 7513 32-Core Processor, 251 GiB of RAM, and two NVIDIA RTX 6000 Ada GPUs, which were instrumental in handling the computational demands of this extensive dataset. Perplexity dropped sharply from approximately 4.99 to 1.19 within the first chunk (completed in two days), indicating efficient early-stage learning (Supplementary Fig. S6). After this initial phase, perplexity stabilized around 1.0 for the remaining chunks, suggesting that the model had effectively captured the underlying chromatin state structure early in the training process.

### 4. Fine-tuning for gene expression level classification

We conducted fine-tuning for each downstream application based on the parameters acquired from the pretraining process. ChromBERT follows the BERT-base architecture, consisting of 12 transformer layers, 12 attention heads, and a hidden size of 384. The intermediate layer size was 1536, and we applied the GELU activation function. The model uses a vocabulary size of 50,630, corresponding to all possible 4-mers derived from chromatin state sequences. The maximum sequence length was set to 512, with dropout probabilities of 0.1 for attention and hidden layers.

We followed the fine-tuning conditions used in DNABERT, applying AdamW as the optimization method with a weight decay of 0.01. Fine-tuning was conducted using a batch size of 32, a learning rate of 2e-4, and a warm-up proportion of 10% of the total training steps. Each fine-tuning task was trained for 10 epochs, with evaluation performed every 100 steps. Model checkpoints were saved every 4000 steps, and gradient accumulation was used to efficiently train on large-scale datasets.

For pairwise binary classification across different gene expression levels, we utilized RNA-seq data from 57 different cell lines provided by the ROADMAP project. A genic region was classified as ‘highly expressed’ if its log-transformed RPKM (Reads Per Kilobase per Million mapped reads) was greater than 5, while genic regions with an RPKM value of 0 were considered ‘non-expressed’. To compare genes with different expression levels, we established five log-transformed RPKM thresholds: 1, 2, 3, 4, and 5. These thresholds were determined based on a preliminary distribution analysis, and they resulted in more balanced gene counts across classes.

For promoter-based classification tasks, ChromBERT was fine-tuned using input sequences of various lengths upstream and downstream of the TSSs. Since BERT distinguishes upstream from downstream context, chromatin state sequences corresponding to genes on the reverse strand were reversed. During the fine-tuning step, we tried tokenization by sliding *N* characters at once (stride *N*), in addition to normal tokenization by sliding one character (stride 1), to accommodate longer sequences (Supplementary Fig S1). Here we tested strides 2 and 3, allowing us to include up to 290 kbp around the TSS (190 kbp upstream and 100 kbp downstream). For each binary classification task, 30,000 and 6,000 genes were randomly selected for the training and test sets, respectively. The model was trained using a maximum sequence length of 512, batch size of 32, learning rate of 2e-4, and training schedule of 10 epochs. The performance was evaluated using F1 score, mean area under the curve (AUC), and accuracy.

### 5. Implementation of a Regression Model for Gene Expression Prediction

The ChromBERT framework was adapted for regression by modifying its processing pipeline to handle continuous RPKM values. Promoter region sequences were paired with their log-transformed RPKM values. A GeneExpressionProcessor, modeled after StsbProcessor, was developed for reading, processing, and tokenizing chromatin state sequences. Since StsbProcessor is designed for sentence-pair regression and requires paired inputs, a custom processor was implemented to streamline data handling, simplify preprocessing, and align with our regression task. This processor was integrated into the task dictionary under the task name “gene_expression”, enabling quantitative gene expression prediction rather than categorical classification. The fine-tuning script was updated accordingly to support regression.

Unlike classification tasks with balanced RPKM thresholds, regression required handling skewed RPKM distributions. The dataset was dominated by low or zero RPKM values, which could bias loss optimization (e.g., Mean Squared Error) and degrade prediction accuracy for high-expression cases. Sequences composed entirely of the “O” state correlated strongly with non-expression, contributing to a higher fraction of zero RPKM values (12.92%) compared to datasets excluding them (8.31%) (Supplementary Fig. S7). To reduce redundancy and bias, these sequences were excluded from the primary dataset.

For gene expression prediction, we fine-tuned a regression model under three TSS-anchored genomic window conditions: 90 kbp upstream and 5 kbp downstream of the TSSs with a stride of 1,190 kb upstream and 5 kb downstream with a stride of 2, and 100 kb upstream and 90 kb downstream with a stride of 2. The model was trained with a maximum sequence length of 512, using a batch size of 32, a learning rate of 2e-5, and a training schedule of 10 epochs. The performance was assessed using Pearson correlation coefficients to measure the consistency between predicted and actual gene expression levels.

### 6. Fine-tuning for cell-type classification

For cell-type classification, we used the cell group annotations defined by the ROADMAP, including “ESC,” “iPSC,” “ES-derived,” “blood-Tcell,” “HSC-Bcell,” “Brain,” “Muscle,” “Heart,” and “Smooth Muscle.” We excluded cell groups with small sample sizes and those with coarse annotations. We used genomic regions of cis-regulatory modules (CRMs) for classification because they are likely to include most functional epigenomic regions. We obtained the reference CRM region lists for genome build hg38 from the SCREEN database 3/17/26 1:55:00 PM and converted them to genomic coordinates for build hg19 using the UCSC LiftOver tool. We removed the CRMs in chromosomes X and Y. The adjacent regions within 200 bp were merged.

Then we extracted the chromatin state sequences from 20-kbp regions around the center of each CRM site for each sample. We merged CRM sequences from all samples for each group. For fine-tuning, we randomly selected 20,000 and 4,000 sequences for training and testing, respectively. Both training and testing data contained an equal number of positive and negative samples corresponding to two cell types. To prevent data leakage due to overlapping genomic regions, training and evaluation datasets were separated by chromosome: chromosomes 1–19 for training and chromosomes 20–22 for evaluation. The model was trained using a batch size of 16, a learning rate of 2e-5, a training schedule of three epochs, and a warm-up proportion of 10% of the total training steps. Model checkpoints were saved every 5,000 steps, and gradient accumulation was applied to enable efficient training on large-scale datasets.

For the cell-type classification using TSSs, we used 20 kbp upstream and 5 kbp downstream of the TSSs with a stride of 1. We used 17,756 expressed genes after excluding non-expressed genes with an RPKM value of 0. To prevent data leakage, we excluded the overlapping regions from the dataset. We then separated the data into chromosomes 1–17 for training and chromosomes 18–22 for evaluation. 30,000 and 5,000 genes were randomly selected for the training and test sets, respectively. The model was trained using a maximum sequence length of 512, the batch size 32, learning rate 2e-5, and training schedule 10 epochs.

### 7. Fine-tuning for 3D genome classification

We used *in situ* Hi-C data of the cell lines GM12878, HMEC, HUVEC, IMR90, and K562 obtained from Rao *et al.* [37]. We downloaded the .hic files for the genome build hg19 with MAPQ ≥ 30 from the Gene Expression Omnibus (GEO) database under accession number GSE63525. The .hic were processed using CustardPy [43] to generate genome-wide Hi-C contact matrices with 25-kbp resolution and the square-root vanilla coverage (VC-SQRT) normalization.

The compartments A and B were identified by performing a principal component analysis (PCA) on the normalized contact matrices, and genomic coordinates were assigned according to the sign of the first eigenvector (PC1). To capture the strength of compartmentalization, the compartments were further classified into four categories based on their corresponding PC1 values: Strong A (PC1 > 75th percentile), Weak A (median < PC1 ≤ 75th percentile), Weak B (25th percentile < PC1 ≤ median), and Strong B (PC1 ≤ 25th percentile). These categories were then segmented into 25-kb genomic bins and used as input data for fine-tuning.

TADs were identified using the Arrowhead function of Juicer [44] at a 25-kb resolution. TAD boundaries were defined as regions located within 25 kb of either end of TADs that exhibited an insulation score [45] of ≤ 0.7, which is a quantitative metric of the boundary strength. Non-TAD boundaries were defined as regions located within TADs that were positioned at least 100 kb away from any TAD boundary and exhibiting an insulation score of > 0.7. The identified TAD boundaries and non-TAD boundaries were merged across five cell types and used for fine-tuning.

For compartment classification, we used the 25 kbp regions with a stride of 1. We used chromosomes 3–22 for training and chromosomes 1 and 2 for evaluation because the later chromosomes show an imbalanced proportion of regions assigned to compartments A and B [37]. We randomly selected 50,000 and 5,000 regions for the training and test sets, respectively. The model was trained using a maximum sequence length of 512, batch size of 32, learning rate of 2e-5, and training schedule of 10 epochs. For TAD boundary classification, we used the 50 kbp regions with a stride of 1 and the 200 kbp regions with a stride of 2 centered around the regions. We used chromosomes 3–22 for training and chromosomes 1 and 2 for evaluation. We randomly selected 10,000 and 2,000 regions for the training and test sets, respectively. The parameter set is the same as that used for the compartment analysis.

### 8. Motif finding

Similarly to the definition of DNA motifs in DNABERT, chromatin state motifs in our study are defined as frequently appearing patterns of consecutive chromatin states that are distinctively present in a certain category during binary classification. We followed the parameter conditions used in DNABERT to detect and filter these motifs. Firstly, we captured regions with high attention scores from the output attention matrix. A threshold was defined as a score higher than the mean attention scores of the matrix and higher than 10 times the minimum score to determine regions where motifs are likely included. These regions were registered as candidates, which were further filtered using a false discovery rate (FDR) threshold of 0.05, based on a hypergeometric test followed by a Benjamini–Hochberg correction. This FDR determined whether the motifs were significantly enriched in one class compared to the other. Finally, motifs were identified as regions that are longer than four chromatin state sequences (equivalent to 800 bps) and appear at least three times exclusively in the target class. While in the case of DNA motifs, final motifs are typically compared with existing DNA motif libraries, we collected chromatin state motifs without such a comparison because there is currently no known library available for this task.

### 9. Motif clustering

To further estimate the representative motif chromatin sequence pattern, we utilized DTW and agglomerative clustering to analyze the chromatin state motifs. Our initial dataset included motifs identified after careful screening for p-values, minimum length, and minimum occurrences within predefined regions of interest, avoiding the merging step to allow a detailed examination of chromatin sequences, which are represented alphabetically. Given the potential for variability in these sequences due to factors such as data quality (e.g., the signal-to-noise ratio of the antibodies) or fluctuations in histone modification peaks, we opted for a more flexible approach to motif identification. DTW was selected for its capability to adjust for sequence length variations, accommodating the intrinsic dynamics of chromatin state sequences.

First, to implement DTW, we converted the chromatin state alphabets (“A” to “O”) into numerical values (1 to 15) to facilitate a standardized analysis. We provide two options for handling this numerical translation of chromatin states by offering a ‘categorical’ parameter, which can be set to ‘True’ or ‘False’. If users set it to ‘True’, the numbers are considered categorical, meaning the states A to B and A to O are equally distant. On the other hand, if it is set to ‘False’, which is the default option, the numbers representing the states are considered numerically; thus, the distance from state A to C is twice that of the distance from state A to B. The result of motif clustering when the ‘categorical’ parameter is set to ‘True’ is as shown in Supplementary Figure S8. Next, Shorter sequences were padded with ‘NaN’ (Not a Number) to match the maximum length dictated by the window size, ensuring consistency across all entries. For DTW calculation, the NaN values in shorter sequences are, by default, converted to the number representing the nearest state according to the ‘ffill’ filling method. Since chromatin state annotations are typically strand-agnostic and assigned without reference to gene orientation, we considered both forward–forward and forward–reverse comparisons to account for potential reverse-orientation patterns in chromatin state motifs. Consequently, the DTW matrix was composed of the minimum score between these two comparisons. Using the ‘tslearn’ Python package, particularly the ‘dtw’ module, we calculated similarity scores between sequences by focusing on the minimum cumulative distances in both directions, allowing for a detailed comparison of motifs.

Following this similarity assessment, we employed agglomerative clustering via the ‘AgglomerativeClustering’ module from the same package, grouping the motifs into clusters based on their similarities. Additionally, we provide a function to generate a dendrogram, which displays the hierarchy and helps users determine the optimal number of clusters when the number of clusters is not predetermined. While the dendrogram was our primary tool for selecting the optimal number of clusters, users are encouraged to consider alternative methods for determining the cluster count. For visualization purposes, we provide two functions: the first displays the UMAP, which reduces the dimensionality of the data for easier interpretation and visualization of the clustering results; the second visualizes the final result of clustering with actual motif entries. This approach facilitates the identification of representative sequential patterns among chromatin states, thereby offering deeper insights into genomic regulatory landscapes.

## Data availability

The pretraining weights of ChromBERT have been submitted to Zenodo (https://doi.org/10.5281/zenodo.15518584).

## Code availability

The source code of ChromBERT is available at https://github.com/caocao0525/ChromBERT.

## Competing interest statement

The authors declare no competing interests.

## Acknowledgements

We thank all the members of the Nakato and Lin laboratories for their helpful discussions. We also thank the IHEC project for allowing us to use reprocessed and harmonized epigenomic data from a large collection of human cell types in this study.

## Authors’ contributions

S.L. developed ChromBERT and performed all analyses in this study. S.L. and J.S. performed a quantitative prediction of gene expression. G.M.O. and R.N. performed the cell-type classification. S.L., Y.N., and R.N. performed the 3D genome classification. S.L. and R.N. performed the binary classification of gene expression. R.N. conceived and designed the study. S.L. and R.N. contributed to the drafting of the manuscript. C.L. and C.Y.C. supervised the deep learning process and provided suggestions for improving the analysis and the manuscript.

## Funding

This work was supported by a Grant-in-Aid for Scientific Research under grant number 23H02466, Grant-in-Aid for Early-Career Scientists under grant number 25K18447, the Japan Agency for Medical Research and Development under grant number JP23gm6310012h0004, and the JST FOREST Program under grant number JPMJFR224Y.

## Competing interests

The authors declare no competing interests.

## Notes

### Competing Interest Statement

The authors have declared no competing interest.

### Summary of Updates

In this revision, we substantially clarified the scope and conceptual positioning of the study. The manuscript has been revised to more clearly define the primary goal as introducing the concept of chromatin state motifs and demonstrating how they can be systematically extracted from chromatin state sequences. We emphasize that ChromBERT is designed to explore how much biological information can be inferred from chromatin state sequences alone, rather than to maximize predictive performance through integration of multiple data modalities. Related sections throughout the manuscript were carefully revised to avoid overstatement and to better articulate the conceptual contribution of this work. We also expanded the downstream evaluation of ChromBERT to further demonstrate its applicability across different biological tasks. In addition to the previously included binary and quantitative gene expression prediction analyses, we incorporated new tasks including cell type classification and three-dimensional genome feature classification. These analyses produced biologically interpretable results and enabled the identification of characteristic chromatin state motifs associated with each task. The pretraining and modeling strategy was also improved to ensure fair and consistent evaluation across tasks. Specifically, the pretraining approach was changed from promoter-based modeling to whole-genome-based modeling. In addition, we introduced a stride-based tokenization scheme to better utilize ChromBERT's ability to process long input sequences. For transcription start site-based prediction tasks, chromatin state sequences corresponding to genes on the reverse strand were reversed so that upstream and downstream regulatory contexts could be properly represented. Finally, we refined the expression modeling framework. Regression analyses indicated that log-transformed expression values produce more stable distributions and improved predictive performance. Based on these observations, the binary classification thresholds were redefined using log-transformed expression values to achieve greater consistency and robustness across analyses.

