## Supplementary figure for "ChromBERT: Uncovering Chromatin State Motifs in the Human Genome Using a BERT-based Approach"

**Supplement Figures**

Figure S1.

Stride-based tokenization of chromatin state sequences. Chromatin state sequences were tokenized using sliding windows with strides of 1–3, reducing sequence length and enabling longer genomic regions to be used as model input.

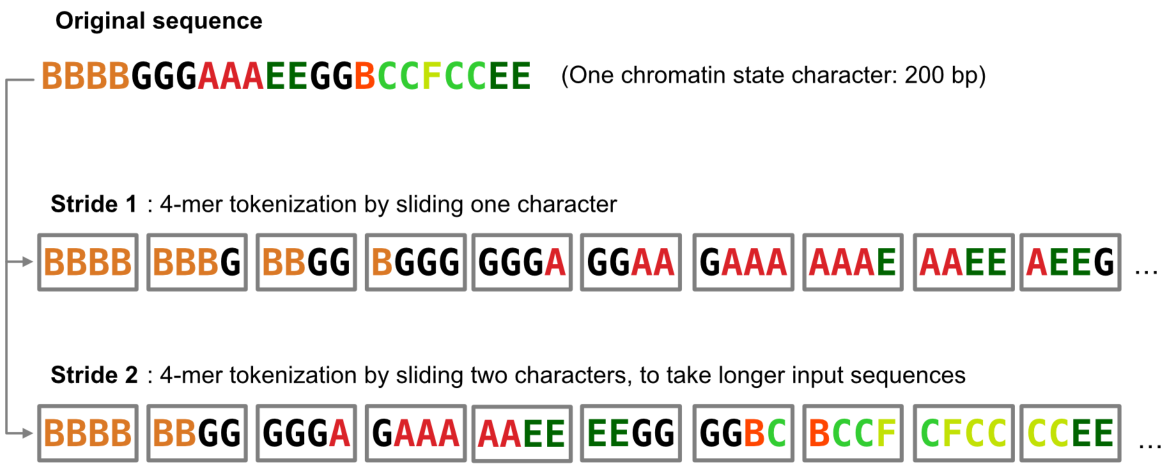

Figure S2.

AUC and accuracy heatmaps corresponding to the upstream/downstream input-region configurations tested in Figure 3(b).

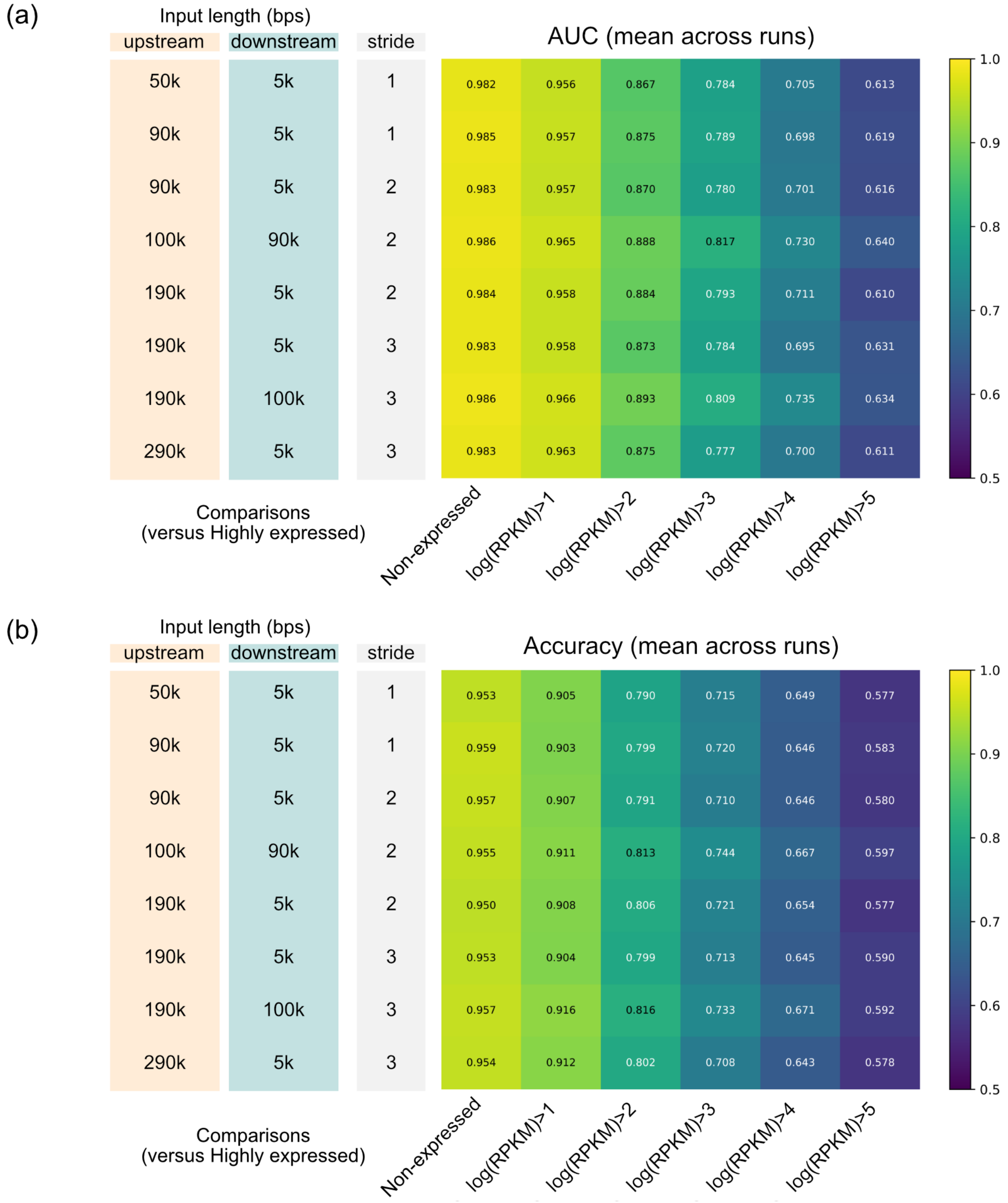

Figure S3.
Line plot representation of all clustered chromatin state motifs. Each line represents a motif, with the x-axis indicating the position within the sequence and the y-axis showing the chromatin state labels.

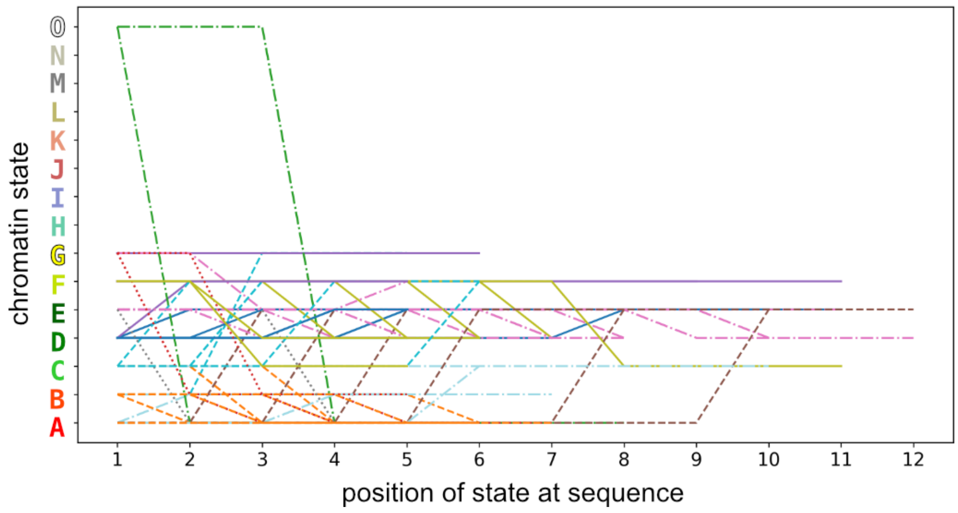

Figure S4.

F1 score heatmap for pairwise binary classification of ROADMAP cell-type groups using chromatin state sequences extracted from TSS-centered windows.

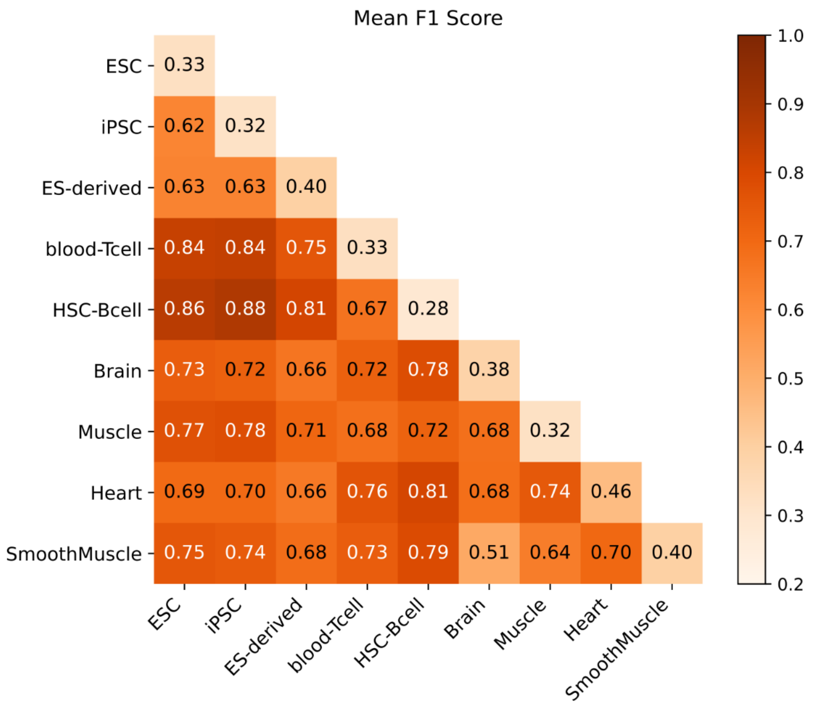

Figure S5.
Examples of chromatin state sequences and their associated RPKM values.

| Sequence | RPKM |
|---|---|
| NNNNNNNNGBAABBBBGGGGGGEEHHEEEEE | 21.19 |
| EEEEEEGAAAAAAAAABBGGGGGFFFFFFFF | 53.977 |
| LMMNNNNNNNNNNNNNNNNNNNNNNNNNNNN | 0.008 |
| GFFFDDDDHHEEEGBBBBAAAAAABBNMMMM | 2.929 |
| EEEEEEEEGGBBBGGNNNNLLGGGGGGGGGG | 0.33 |
| NNNNNNNNNNNNMMMMMLKKKKLLEEEEEEE | 0.985 |
| EEEEEEEEEEEEEEEEEEEEEEEEEEEEEEE | 0.04 |
| DDDDDDDEEEEEEEEAAAAAAAAACCFFFEE | 42.323 |
| EEEEAAAAAAAAACCFFFEEEEEEEEEEEEE | 12.099 |
| NNNNNNNNNNNNNNNNLLLKJJLLMMMMMEE | 1.181 |
| EEEEEEEEEEEEEEBBBBAAAAAAAKOOOOO | 12.686 |
| OOOOOOOOOOOOOOOOOOOOOOOONNNNNNN | 0.726 |
| EEEGGGGGGGGGGGBBBAAAAAAAAAABBCF | 4.375 |
| OOOOOOOOOOOOOOOOOOOOOOOOOOOOOOO | 0.018 |
| OOOOOOOONNNNNNNNNNNNMMMLLMLKKJM | 0.006 |
| OOOOOOAAAAAAAAAAEEDDDDDDDDDDDDD | 83.806 |
| EEEEEEGGGGGGAAAAAAAAAAAKKKGGGGO | 26.401 |
| GLLMMMMMMMMJJKJKKMMNNNNNNNNNNNN | 2.397 |
| OOOOOOOOGGBBBAAAAAAAAAAAAABEEEE | 19.179 |
| FFFFFFFFFCCCCCCAAAAAAAAAABGGGGG | 377.137 |
| GGGGGGGGGOOOOOOOOOOOOOOOOOOOOOO | 0.002 |
| OOOOOOOOOOOOOOOOOOOOOOOOOOOOOOO | 0.011 |
| OOOOOOOOOOOOOOOOOOOOOOOOOOOOOOO | 0.02 |
| AAAAKKLLLLLLLLLLMMJJJJJKLMMMMMM | 0.033 |
| EEEGGGEEEGGGGGOOOOOOOOOOOOOOOOO | 0.017 |
| GGBBBAAAAAAAAAABBGEEEEDDDDEEEEE | 19.201 |
| JJJJLLMMMMMLLLLGGGGGGGGGGGEEEEE | 0.26 |
| IIIIIIIIIIIIIIIIIIIIIIIIIIIIIII | 0.029 |
| IIIIIIIIIIIIIIIIIIIIIIIIIIIIIII | 0.004 |
| LLLKKKJJJJJJJJJJJJJJLLLLLLLLLLL | 0.05 |
| OOOOGGGGGBAAAAAABBBGGEEEEEEEEEE | 3.991 |
| EEEEEEGGGGGGGGGGBBAAAAAKKLLLLGG | 3.609 |
| NNNNNBKAAAAAABBBEEEEEEDDDDDDDDD | 13.04 |
| DDDDDDDDDEEEEEEEBBAAAAJJKKKKKJJ | 4.554 |
| JJKKKKKJJJJJJKJLKMMMMMMMMMMMMMM | 1.257 |
| NNNNNNNNNNNNNNNNNNNNNNNNNNNNNNN | 0.014 |

Figure S6.

Results of pretraining for whole genome using IHEC dataset (1699 different epigenomes).

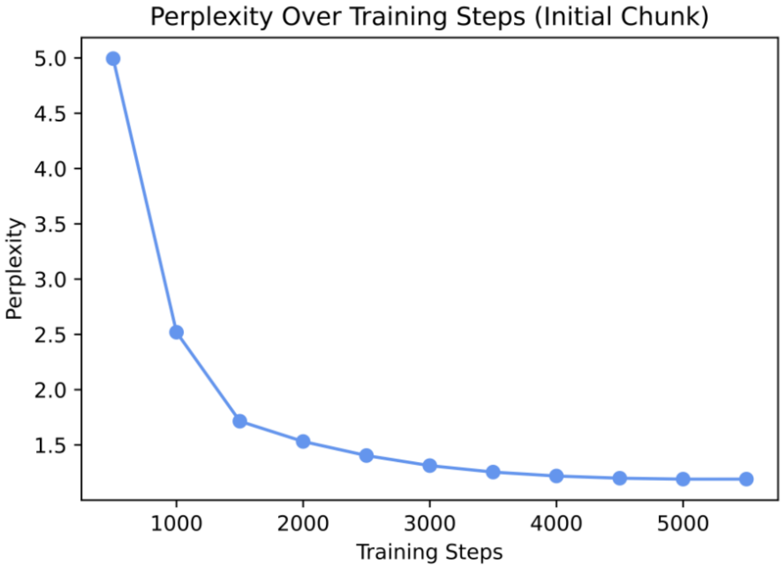

Figure S7.
Comparison of RPKM distributions in promoter regions between datasets with and without entire ‘O’-state sequences.

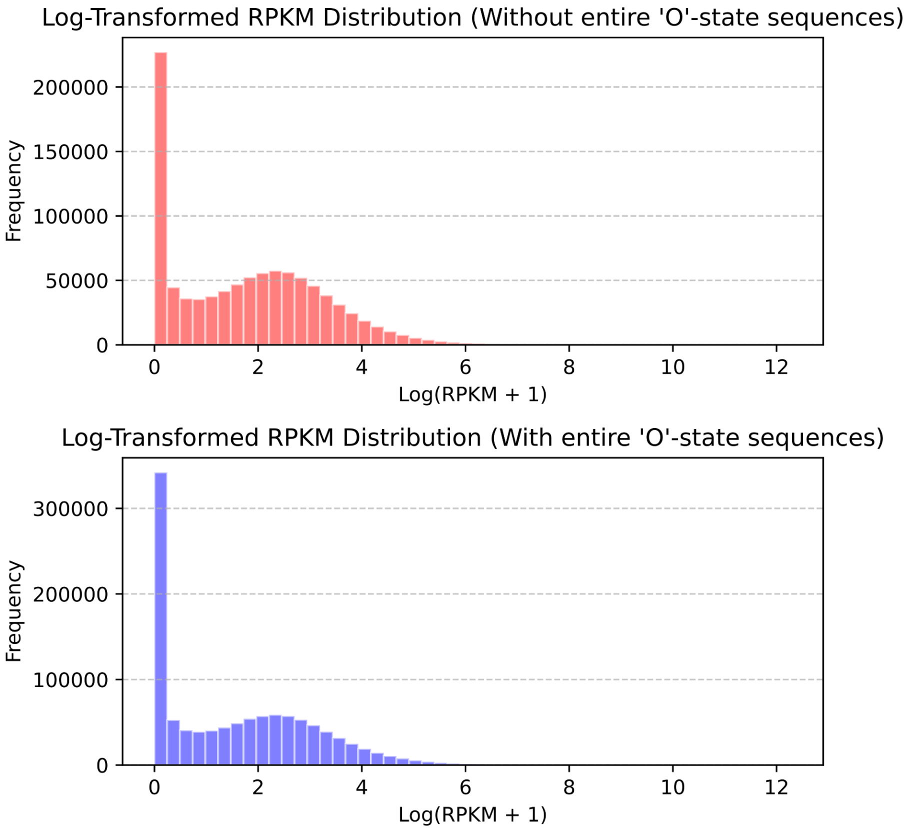

Figure S8.

Motif clustering result when the parameter ‘categorical’ is set to ‘True’ (a) UMAP representation (b) Visualization of all clustered motifs.

(a)

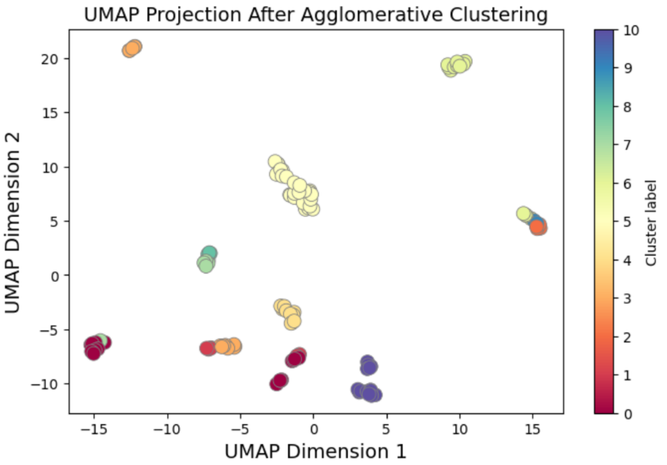


(b)

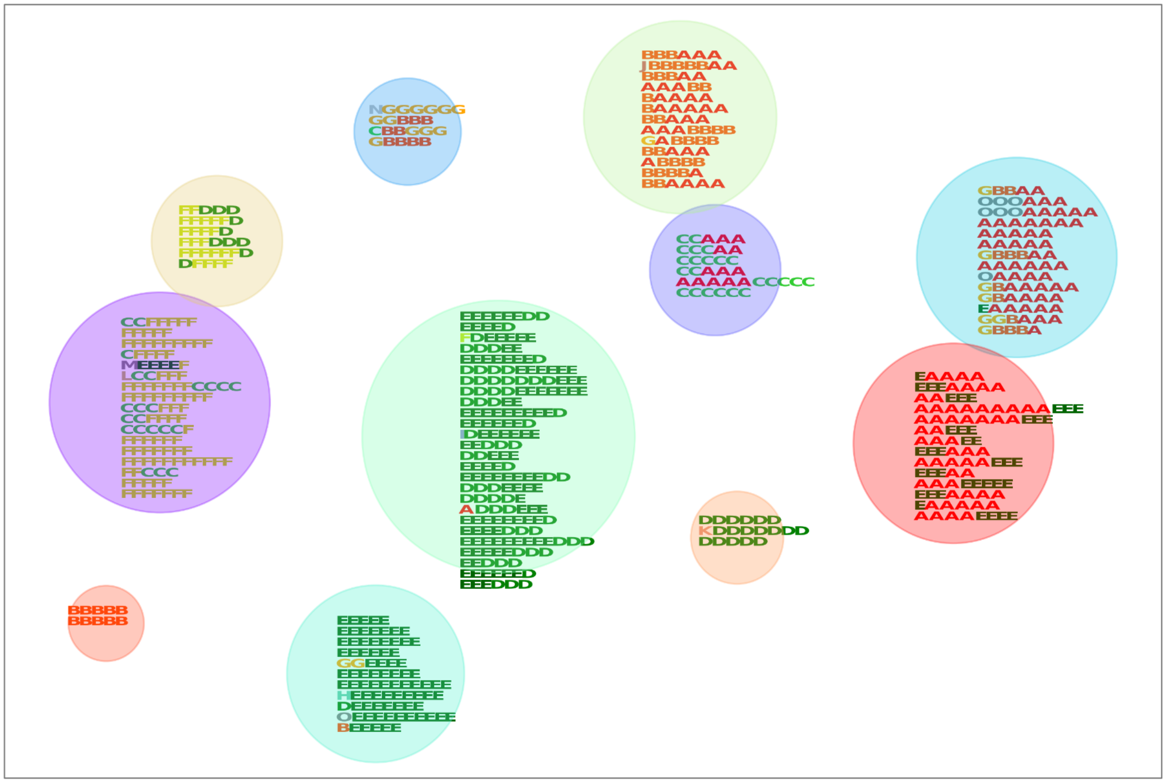
